# Tracking extracellular vesicle (EV) cargo as a platform for studying EVomics, signaling, and targeting *in vivo*

**DOI:** 10.1101/2021.09.23.461577

**Authors:** Inna A. Nikonorova, Juan Wang, Alexander L. Cope, Peter Tilton, Kaiden M. Power, Jonathon D. Walsh, Jyothi S. Akella, Amber R. Krauchunas, Premal Shah, Maureen M. Barr

## Abstract

Extracellular vesicle (EV)-based signaling is a challenge to study, due to EV small size, heterogeneity, and limited information on cargo content *in vivo*. We present *Caenorhabditis elegans* as a discovery platform that allows single EV tracking from source to target tissue in living animals. We enriched ciliary EVs using GFP-tagged PKD-2 cargo followed by mass spectrometry analysis to identify 2,888 cargo candidates. By integrating our dataset with single-cell transcriptomic data, we identified EV cargo produced by individual neurons and other cell and tissue types. A single cilium produces multiple EVs with distinct protein content. Ciliary EVs carry nucleic acid binding proteins. We observed transfer of EV cargo from the male reproductive tract to the hermaphrodite uterus during mating, a direct demonstration of animal-to-animal EV targeting.

**One-Sentence Summary:** Here we present a discovery platform for studying animal extracellular vesicle composition and biogenesis.

## Main Text

Extracellular vesicles (EVs) represent an ancient and conserved form of distant inter-cellular and inter-organismal communication in different physiological and pathological states. EVs carry bioactive cargo to direct development and differentiation, initiate synapse formation in neurons, modulate maturation and fertilization of gametes [1–4]. EVs also may spread toxic cargo, such as unfolded proteins in neurodegenerative diseases or signaling molecules to establish a metastatic niche for cancer cells [5,6]. While EVs are of profound medical importance, the field lacks a basic understanding of how EVs form, what cargo is packaged in different types of EVs originating from same or different cell types, and how different cargoes influence the range of EV targeting and bioactivities.

Our study focuses on ciliary EVs that bud from the membrane of cilia, the cellular antennae that receive and transmit signals for intercellular communication. Disrupted ciliary EV signaling is likely an important driver of the pathophysiology of many ciliopathies, such as polycystic kidney disease, retinal degeneration, and others [7]. Ciliary EV biogenesis is an evolutionarily conserved process for which main discoveries were pioneered in the green algae *Chlamydomonas reinhardtii* and free-living nematode *Caenorhabditis elegans* [8,9]. Nematodes use EVs to communicate with mating partners and to modulate and evade host immune responses via specific EV cargo [8,10–15]. Here we used *C. elegans* as a discovery platform to conduct a large-scale isolation and proteomic profiling of environmentally released EVs.

### PKD-2, an evolutionarily conserved ciliary EV cargo, enables EV visualization and tracking during biochemical enrichment

Our strategy for biochemical EV enrichment relied on the use of a *C. elegans* strain expressing GFP-labeled polycystin-2 (*pkd-2::gfp*), which encodes an evolutionarily conserved cargo of ciliary EVs [8,11]. EVs carrying PKD-2 are shed from the cilia of male-specific sensory neurons (<u>ce</u>phalic <u>m</u>ale CEM and tail <u>r</u>ay RnB and <u>h</u>ook HOB neurons) into the environment. PKD-2-carrying EVs function in inter-organismal communication by changing male locomotion. Males transfer PKD-2::GFP carrying EVs to vulvae of their mating partners [11,12]. In addition to the PKD-2-expressing sensory neurons, *C. elegans* has a set of sex-shared ciliated neurons whose cilia are exposed and release EVs to the environment – six <u>i</u>nner <u>l</u>abial type <u>2</u> (IL2) neurons (Figure 1A). The male specific PKD-2 expressing neurons and IL2s are collectively termed EV-releasing sensory neurons (EVNs) for their ability to release ciliary EVs with unique and/or common cargo, whereas ciliary EV shedding has not been directly observed from the sex-shared major sensory organs in the head (amphids) and tail (phasmids). Since PKD-2 is a male-specific EV cargo, we used a strain with high incidence of males (*him-5*) and cultured the mixed-sex population on standard bacterial lawns of *E. coli* OP50 favorable for mating to promote the release of signaling ciliary PKD-2 EVs.

**Figure 1.**
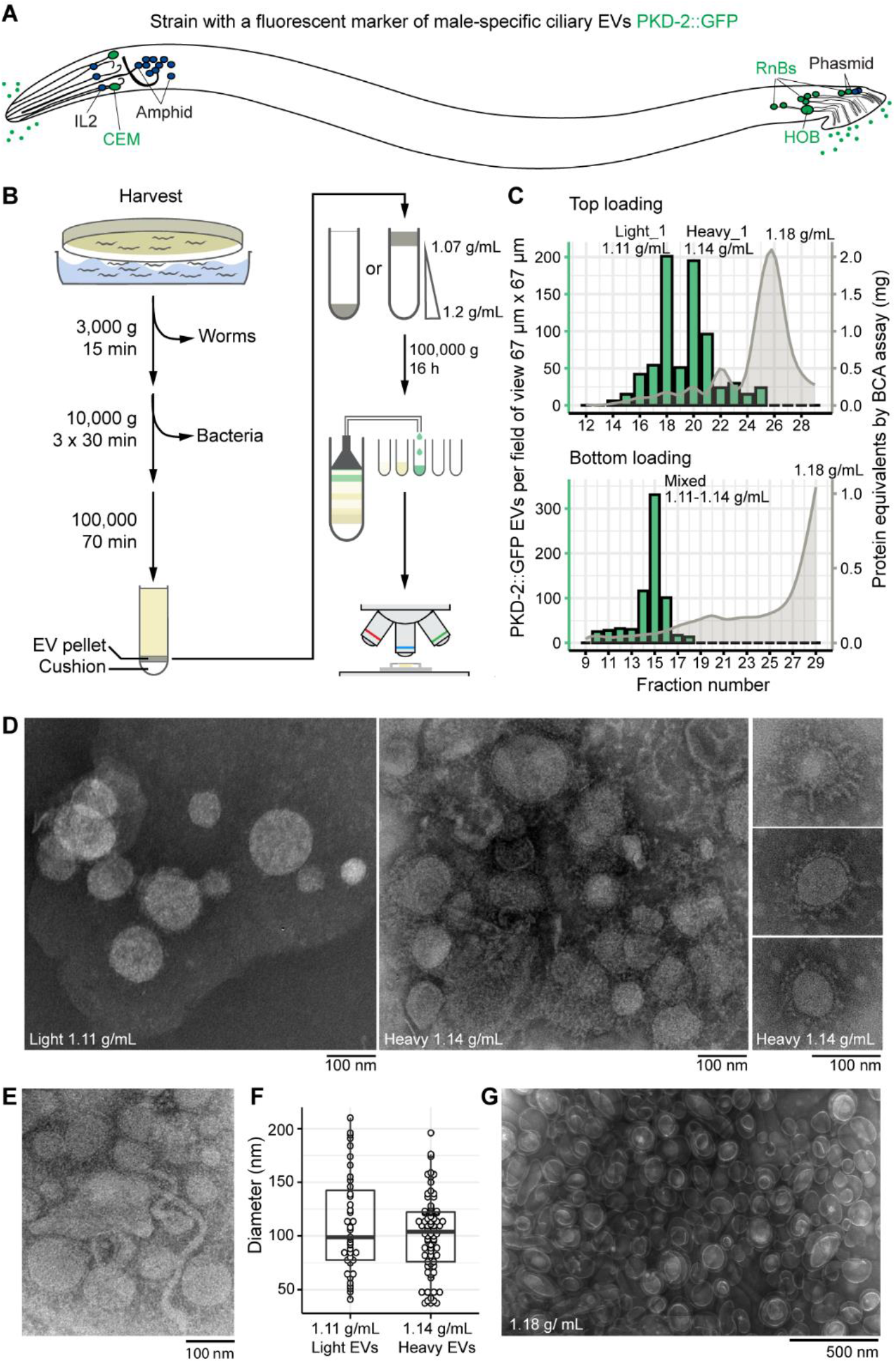
Buoyant density centrifugation enriches for ciliary PKD-2::GFP EVs more than 100-fold and resolves them into two populations. (**A**) Location of environmentally exposed cilia of sensory neurons. Male-specific cilia are labeled in green. Neurons: CEM – cephalic male, IL2 – inner labial type 2, HOB – hook type B, RnBs – ray neurons type B. (**B**) Schematic of the EV enrichment workflow. (**C**) Combination charts showing number of the PKD-2::GFP EVs and the amount of protein in the collected fractions. Centrifugation of the top-loaded sample resolved PKD-2 EVs in two types with densities 1.11 g/ml (light) and 1.14 g/mL (heavy), respectively. Bottom-loaded samples failed to resolve the PKD-2 EV subtypes but reached a similar level of enrichment (more than 100-fold). (**D**) Transmission electron microscopy (TEM) of the negatively stained PKD-2 EV enriched fractions revealed presence of vesicles with coatings. (**E**) Tubular structures of 17-20 nm in diameter reminiscent of the stem of a budding vesicle. (**F**) Diameters of EVs recovered from the light and heavy fractions fall in the range of 50 – 200 nm as measured by TEM. (**G**) EVs from the fraction with most protein, presumably bacterial OMVs.

### Ciliary EV enrichment method

EVs were collected using ultracentrifugation after serial enrichment by several differential centrifugation steps. The resulting crude EV mixture was then resolved using buoyant density centrifugation and high-resolution fractionation (Figure 1B). Collected fractions were profiled to determine their density, protein concentration, and the quantity of PKD-2::GFP EVs using Airyscan confocal imaging (Figure 1C, S1A).

In accordance with best practices for EV isolation [16], we tested two protocols of loading EVs for buoyant density centrifugation, top- and bottom-loading for top-down (pelleting) and bottom-up (flotation) gradient equilibration, respectively (Figure 1B). The top-loading method resolved PKD-2::GFP EVs into two sub-populations of 1.11 g/mL (termed “light”) and of 1.14 g/mL (termed “heavy”) densities. The bottom loading resulted in all PKD-2 EVs migrating as a single band that contained both light and heavy PKD-2 EVs (termed “mixed”). Analysis of protein distribution across gradients for both loading approaches showed that 99% of total protein equilibrated at density of 1.18 g/mL and was not associated with the labeled EVs (Figure 1C), indicating that we achieved at least 100-fold enrichment for ciliary PKD-2::GFP EVs.

We confirmed the presence of EVs in the fractions-of-interest using transmission electron microscopy (Figure 1D-E). EVs from the PKD-2::GFP-enriched fractions ranged in size from 50 nm to 220 nm in diameter (Figure 1F), suggesting heterogeneity of the isolated environmental EVome. In addition to intact EVs, we also observed tubular structures reminiscent of the budding vesicle stems [17] (Figure 1E), indicating that our EV preparations contained membrane-sculpting proteins participating in the formation of EVs.

For rigor and reproducibility, we isolated EVs in two more biological replicates using the top-loading method. Each time, PKD-2::GFP-carrying EVs resolved into light and heavy sub-populations. Electrophoresis of proteins isolated from collected EVs revealed greater heterogeneity of the PKD-2 enriched fractions as compared to the fraction with the most protein (1.18 g/ml), where the predominant protein was 35-37 kD (Figure S1B). This size range and density of the band corresponded to bacterial outer membrane proteins (OMPs) [18,19], the major constituents of the outer membrane vesicles (OMVs) released by the *E. coli* culture lawn. OMPs were also present in the PKD-2::GFP enriched fractions, with more prominent presence in the heavy fraction. To increase the depth of protein identification in the subsequent mass spectrometry, we chose not to combine technical replicates (parts of the same biological sample that were run in different density gradient tubes) and treated each as separate samples (Figure S1C-S1E). In addition, we excluded the OMP band prior to mass spectrometry to enrich for the presence of *C. elegans* proteins (Figure S1F). In total, we used 14 samples for protein identification analysis: one mixed fraction from the bottom-loading approach and three top-loaded biological replicates represented by one, two, and three technical replicates, respectively (Figure S1E-S1F).

### Protein identification in the fractions enriched with PKD-2::GFP EVs

In total, we obtained 147,132 spectral counts that mapped to the *C. elegans* proteome, with 21,254 being unique protein identifiers for 2,888 *C. elegans* proteins and 1,535 peptides that mapped to more than one *C. elegans* protein (Table S1). Pairwise comparison using Spearman correlation analysis showed reproducibility of our approach. Correlation coefficients ranged within 0.72 – 0.88 between technical replicates and 0.43 – 0.76 between biological replicates (Figure S2A) and 60% of the EV cargo candidates were identified in at least 2 out of 3 biological replicates (Figure S2B). Hierarchical clustering analysis segregated all the light fractions from all the heavy ones (Figure S2C), consistent with the idea that the light and heavy EVs might carry unique sets of proteins and/or different amounts of the same proteins. Qualitatively 58% of identified proteins were common between light and heavy fractions, whereas the mixed fraction shared 95% of its proteins with the light and heavy fractions (Figure S2D) consistent with presence of both light and heavy EVs.

Light EV fraction was enriched in proteins involved in biosynthesis of amino acids, glycolysis, one-carbon metabolism, nucleocytoplasmic transport, and pentose phosphate pathway, and other carbon metabolic processes (Benjamini-Hochberg adjusted p-value < 0.05). Heavy EV fraction was enriched in gene products involved in oxidative phosphorylation, regulators of ER-associated protein processing and export, glycerophospholipid metabolism, N-glycan biosynthesis, and biosynthesis of unsaturated fatty acids (Benjamini-Hochberg adjusted p-value < 0.05). Overall, the identified EVome was enriched in proteins associated with endocytosis, lysosome, proteasome, phagosome, ribosome, SNARE machinery, TCA cycle, and glutathione metabolism (Figure 2A, Table S2).

**Figure 2.**
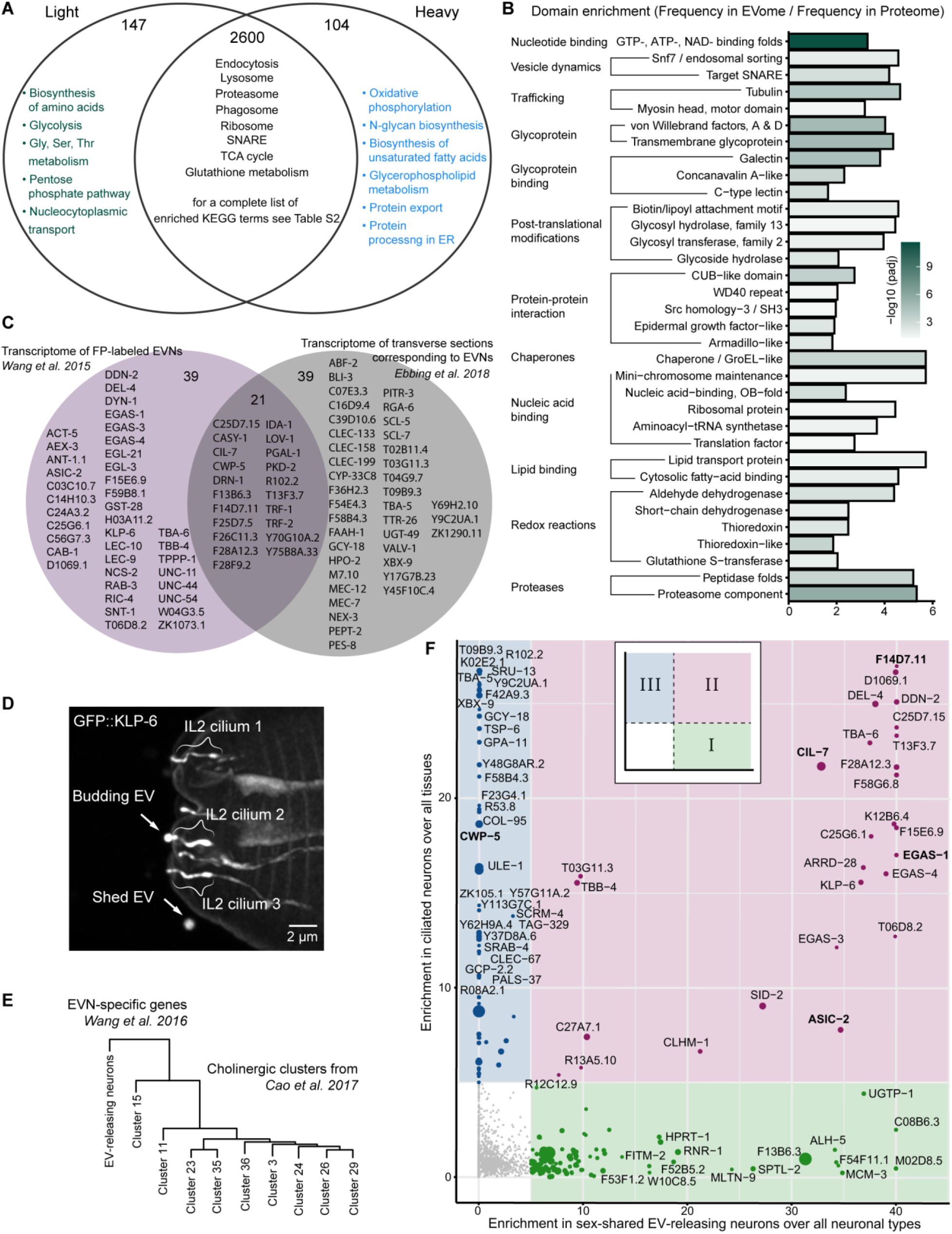
Identified EV proteome is enriched in nucleic acid binding proteins and ciliary EV cargo. (**A**) KEGG categories of pathway analysis that were enriched in the light and heavy EV proteomes. (**B**) InterPro domains enriched in the identified EV proteome (for the full list see Table S2). (**C**) Identified EVome contained proteins encoded by previously identified transcripts specific to EV-releasing neurons (EVNs). The Venn diagram shows EV cargo candidates that overlap with two transcriptomic datasets [10,31]. (**D**) GFP::KLP-6 EV shedding from the IL2 cilia of a wild-type hermaphrodite, orthogonal projection. GFP::KLP-6 was observed in EVs released outside the animal. (**E**) The transcriptome of EV-releasing neurons is most similar to the Cholinergic cluster 15 from [37] likely representing the IL2 neurons. (**F**) EV cargo candidates were categorized into three groups based on their relative enrichment in the IL2 sex-shared EV-releasing neurons (x-axis) and in the ciliated neurons (y-axis). Three categories are indicated by colors. Previously validated ciliary EV cargoes are bolded.

Of 2,888 identified EV cargo candidates, 2,489 were annotated with InterPro domains. Significant enrichment was established for proteins participating in nucleotide binding, vesicular dynamics, trafficking, glycoproteins, glycoprotein binding proteins, nucleic acid binding, post-translational modifiers, chaperones, redox proteins, proteases, and many proteins with domains known to play a role in protein-protein interactions in signaling cascades (Figure 2B, Table S2). Moreover, specific enrichment was observed in proteins carrying Snf7-like domains, major constituents of the endosomal sorting complex required for transport (ESCRT). Specifically, the Snf7-like domains sculpt membrane bilayers into filaments during vesicle budding [17]. Their enrichment in our EV preparations is consistent with observations of the tubular structures by TEM (Figure 1E). For the full list of the identified ESCRT-associated machinery and its human orthologs see Table S2. Role of the ESCRT machinery in ciliary EV biogenesis is suggested by studies in zebrafish [20,21] and *Chlamydomonas* [22,23], but remains largely unexplored.

### Integration with cell-specific transcriptomic datasets

Because the identified EV proteome contained 13% of the entire *C. elegans* proteome, we reasoned that the captured EV cargo candidates likely originated from multiple cellular sources and not solely from the ciliated EV-releasing neurons (EVNs). Nematodes produce EVs from the plasma membrane of embryonic cells [24], epithelial cells and cuticle [25], reproductive tract [26,27], nervous system [11,28,29], and intestine [30]. In these reports, compelling evidence for tissue-specific EV biogenesis was obtained using electron microscopy, whereas EV cargo composition remains unknown.

To identify EV cargo originating from the EVNs, we used cell-specific transcriptomic datasets reporting on EVN-enriched transcripts [10,31]. We found 99 gene products (Figure 2C, Table S3) in our EVome that are also enriched in EVNs, including previously identified and confirmed EV cargo such as the polycystins PKD-2 and LOV-1 (a homolog of human PKD1), a myristoylated coiled-coil protein CIL-7 required for environmental EV biogenesis [32], CWP-5 (a protein <u>c</u>oexpressed <u>w</u>ith <u>p</u>olycystins), acid-sensing sodium channels ASIC-2 and EGAS-1, and a transmembrane cysteine-rich protein F14D7.11 that belongs to a family of proteins with CYSTM-like domains presumably involved in stress-resistance mechanisms [33]. The presence of these EVN-specific cargo in our EV preparations indicates high degree of enrichment for ciliary EVs given the fact that EVNs occupy less than 0.01% of the whole-body volume [34] and are represented by 27 cells in an adult male and just 6 cells in the hermaphrodite out of almost 1000 somatic cells and 2000 germ cells per each animal.

Our dataset also contained EVN-specific ciliary proteins that were previously not observed in the process of shedding from cilia. These include a TGFβ-like domain-containing protein F28A12.3, an ortholog of human ciliary protein CEP104 – T06D8.2, a tetraspanin-like protein T13F3.7, and mucin-domain containing proteins (F25D7.5, F26C11.3, F59A6.3, and Y70C10A.2). Ciliary kinesin-3 KLP-6, previously known as an EV cargo of mutants lacking EVN-specific tubulin TBA-6 [35], was also found in our EV preparations. Re-evaluation of the KLP-6 reporter in the wild-type animals revealed that KLP-6 was shed from the cilia of the IL2 neurons to the environment in the form of ectosomes that bud from ciliary tips (Figure 2D), thereby providing experimental validation for KLP-6 as a member of the EVome. The KLP-6 mammalian homolog KIF13B undergoes EV-release from the ciliary tip in immortalized human retinal pigment epithelial (hTERT-RPE1) cells [36], indicating that cargo of our ciliary EVome are conserved.

Comparison to cell-specific transcriptomic datasets captures only proteins that are enriched in the EVNs and depleted from other tissues. Thus, this comparison does not identify cargo produced by both EVNs and larger tissues, such as the intestine or reproductive tract, for vesicular trafficking and EV biogenesis. To overcome this limitation, we developed a relative scoring approach for EV cargo candidates where their expression in a cell-of-interest was compared to their mean expression value in the corresponding group of functionally related cells (e.g., EVNs compared to all neurons taken together). The relative scoring was made possible by single-cell transcriptomic data obtained from profiling hermaphrodites at the 2^nd^ larval stage L2 [37]. Due to the lack of such data for *C. elegans* males, we used the sex-shared IL2 EVNs as a proxy for the male-specific EVNs, because of their similarities in transcriptional signatures including expression of components of the ciliary EV biogenesis machinery [10]. The IL2 cholinergic neurons were not explicitly identified in the study of Cao *et al* [37], however, all cholinergic neurons of *C. elegans* were segregated into nine types. To determine which of the nine represented the IL2 neurons, we compared their transcriptional signatures to the EVN-specific transcriptome [10] and established that Cholinergic cluster 15 was most related to the EVNs (Figure 2E). We then scored our EV cargoes using two metrics: (x-axis) relative enrichment in the IL2 neurons over mean expression in all neuronal types and (y-axis) relative enrichment in ciliated sensory neurons over mean expression in all tissues (Figure 2F). Scoring EV cargo candidates using these two metrics segregated EV cargo into three categories-of-interest: (i) enriched in IL2 neurons, (ii) enriched in both IL2 and other ciliated neurons, and (iii) enriched in ciliated sensory neurons but excluded from IL2 neurons (Figure 2F). We proceeded with experimental validation of novel EV cargo candidates in all three categories defined by an arbitrary enrichment ratio of more than 5-fold (Figure S3A).

### Cilia produce EVs that carry nucleic acid-binding proteins SID-2, ENPP-1, and MCM-3

SID-2 (<u>s</u>ystemic RNA <u>i</u>nterference <u>d</u>efective) is a double stranded RNA (dsRNA) transporter required for environmental RNAi in *C. elegans*. SID-2 internalizes ingested dsRNA from the intestinal lumen, the first step in the systemic silencing of gene expression [38–40]. The native function of SID-2 remains unknown. Intrigued by this and the abundance of SID-2 in our EV preparations, we examined SID-2 endogenous expression and protein localization patterns using C-terminal mScarlet fluorescent tag, engineered by CRISPR/Cas9-mediated insertion into the *sid-2*-coding region in the genome (Figure S3B). The resulting SID-2::mScarlet reporter was co-expressed with PKD-2::GFP in male-specific EVNs, as well as in intestine and excretory system in both males and hermaphrodites (Figure 3A, S3D-E). SID-2::mScarlet was abundant on the apical surface of the intestine, consistent with its known role in endocytosis od dsRNA (Figure S3C) [38–40].

**Figure 3.**
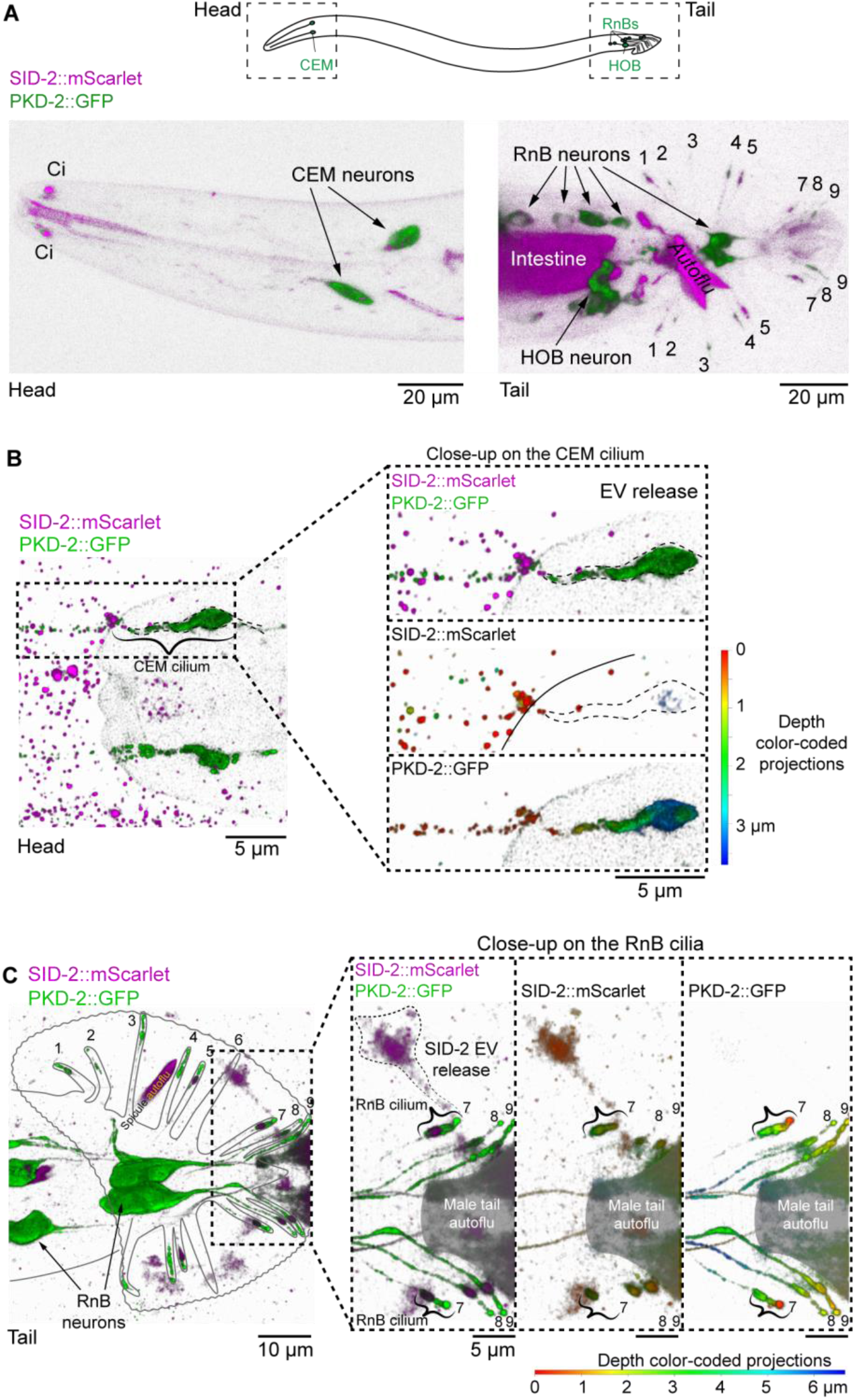
Nucleic acid binding transporter SID-2 is specifically enriched in the male-specific ciliated EV-releasing neurons (EVNs) and is shed in the form of EVs to the environment. (**A-C**) SID-2::mScarlet and PKD-2::GFP reporter presence in the head and tail of an adult male. Zoomed insets in (B) and (C) show ciliary localization and EV release. Areas of intense inherent autofluorescence are shaded with gray and labeled as autoflu. Ci-cilia, CEM – cephalic male neuron, HOB – hook neuron type B, RnB – Ray neuron type B. Merged images are shown as 3D projections; individual channels are presented as depth color-coded projections with minimal labeling where colocalized presence is indicated by the same color (such as SID-2 and PKD-2 presence within the CEM cilium in (B) is indicated with blue which corresponds to ~3.5 μm distance deep into the acquired Z-stack). For individual channels of (A) see Figure S3.

In male-specific EVNs, SID-2::mScarlet was abundant at the ciliary base regions, suggesting that its primary function in these neurons is associated with the cilia. We also observed SID-2::mScarlet being environmentally released in the form of ciliary EVs (Figure 3B-C). Because SID-2 is a transmembrane RNA-binding protein, its presence in ciliary EVs suggests that ciliary EVs might be capable of transporting RNA. To our knowledge this is the first report of an RNA-binding protein as a cargo of ciliary EVs, suggesting that *C. elegans* cilia possess molecular machinery for extracellular transport of RNA.

ENPP-1 (C27A7.1), an ortholog of human <u>e</u>cto<u>n</u>ucleotide <u>p</u>yrophosphatase <u>p</u>hosphodiesterases (ENPPs), is another nucleic acid binding candidate cargo that co-segregated with known EVN-specific EV cargo (Figure 2F). ENPPs are esterases that hydrolyze phosphodiester linkages in either nucleotides (e.g. ENPP1) or lysophospholipids (e.g. ENPP2, aka Autotaxin) [41]. Comparative analysis of C27A7.1 to mammalian ENPP-1 and ENPP-2 showed that C27A7.1 possesses sequence determinants of substrate preference for nucleotides, and not for lysophospholipids [42] (Figure S4A). Hence, we referred to C27A7.1 as ENPP-1 and generated an ENPP-1::mScarlet transgenic translational reporter.

ENPP-1::mScarlet showed a speckled presence in the EVNs, with specific accumulations in the cilia of the CEM and shared IL2 neurons in the head and ray RnB neurons in the male tail (Figure 4A-B, S4B-D). The ENPP-1::mScarlet was released in environmental EVs from both CEMs and IL2s (Figure 4A-B, Movie S1). ENPP-1::mScarlet EV shedding from IL2s was qualitatively different from CEM EV shedding: the tips of IL2 cilia occasionally shed large ENPP-1 and KLP-6 carrying EVs via ectocytosis (Figure 2D), whereas CEMs abundantly shed smaller EVs. Taken together our data establish the cilium as a cellular location for ENPP-1 EV shedding. Consistent with a role in cilia, mammalian ENPP1 loss-of-function manifests signs of ciliopathies, such as accelerated ossification, hearing impairment, altered insulin metabolism and immune response [43–46].

**Figure 4.**
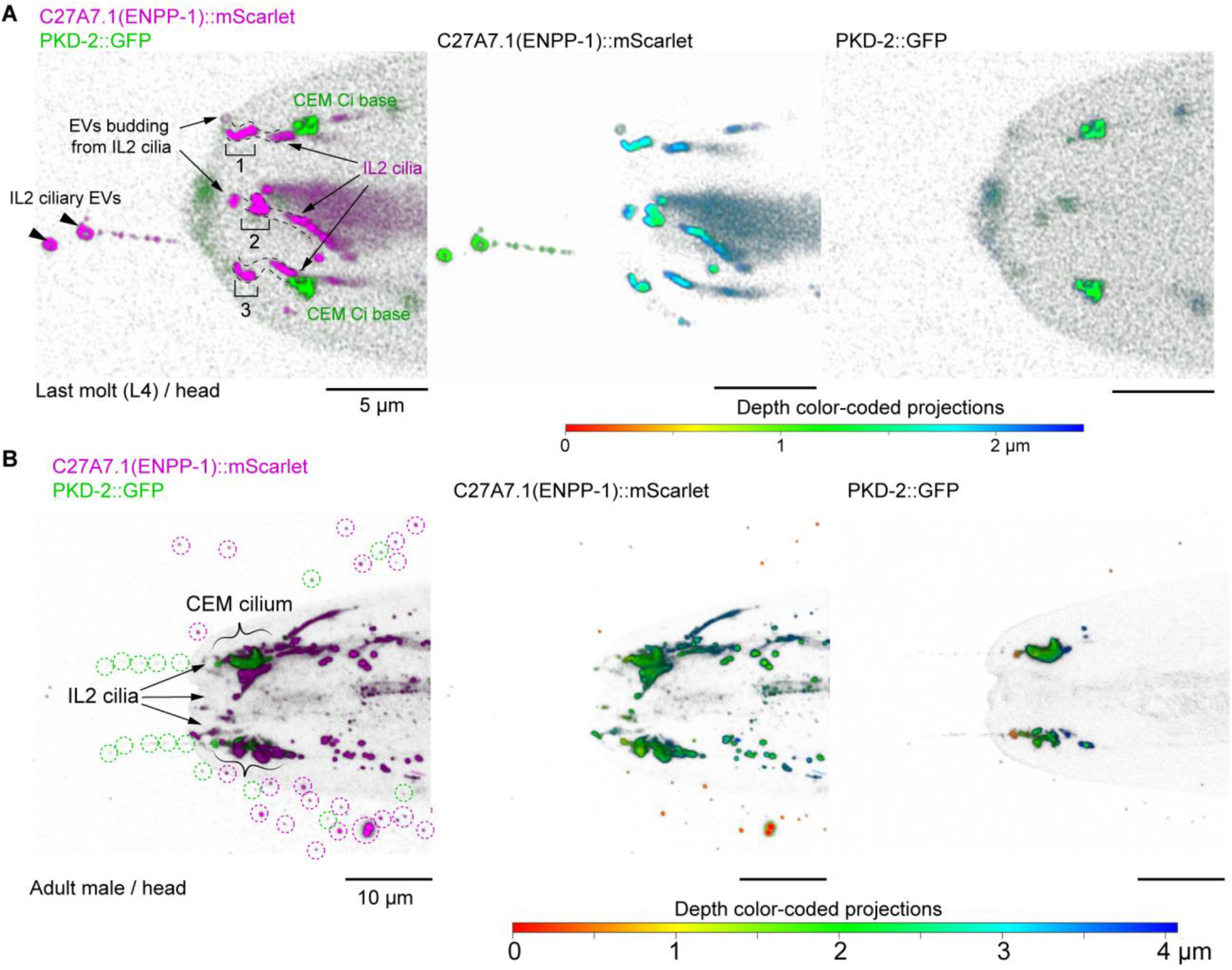
Phosphodiesterase ENPP-1 is a ciliary EV cargo. Ciliary localization of ENPP-1::mScarlet in the sensory neurons of the head: IL2 (**A**) and CEM (**B**). IL2 EVs are indicated by arrowheads, CEM EVs are shown by dashed circles. See movie S1 for time-lapse imaging of ENPP-1::mScarlet ectocytosis from IL2 cilia. PKD-2::GFP served as a marker of the CEM cilia and EVs.

To validate the IL2-enriched category of EV cargo candidates (Category I, Figure 2F), we focused on another nucleic acid binding protein, MCM-3 (<u>m</u>ini<u>c</u>hromosome <u>m</u>aintenance complex component 3 licensing factor) and generated transgenic animals expressing MCM-3::mScarlet driven by its native promoter. Strikingly, MCM-3::mScarlet was specifically enriched in the cytoplasm of all EVNs in adult animals and in their cilia and co-localized with PKD-2::GFP (Figure 5A, S5A-B). MCM-3::mScarlet was released from the cilia in the form of EVs. MCM-3::mScarlet was also expressed in the developing embryo where it showed nuclear localization consistent with the known function of the MCM proteins in DNA replication licensing and DNA helicase activity of the MCM-2-7 complex [47] (Figure S5C).

**Figure 5.**
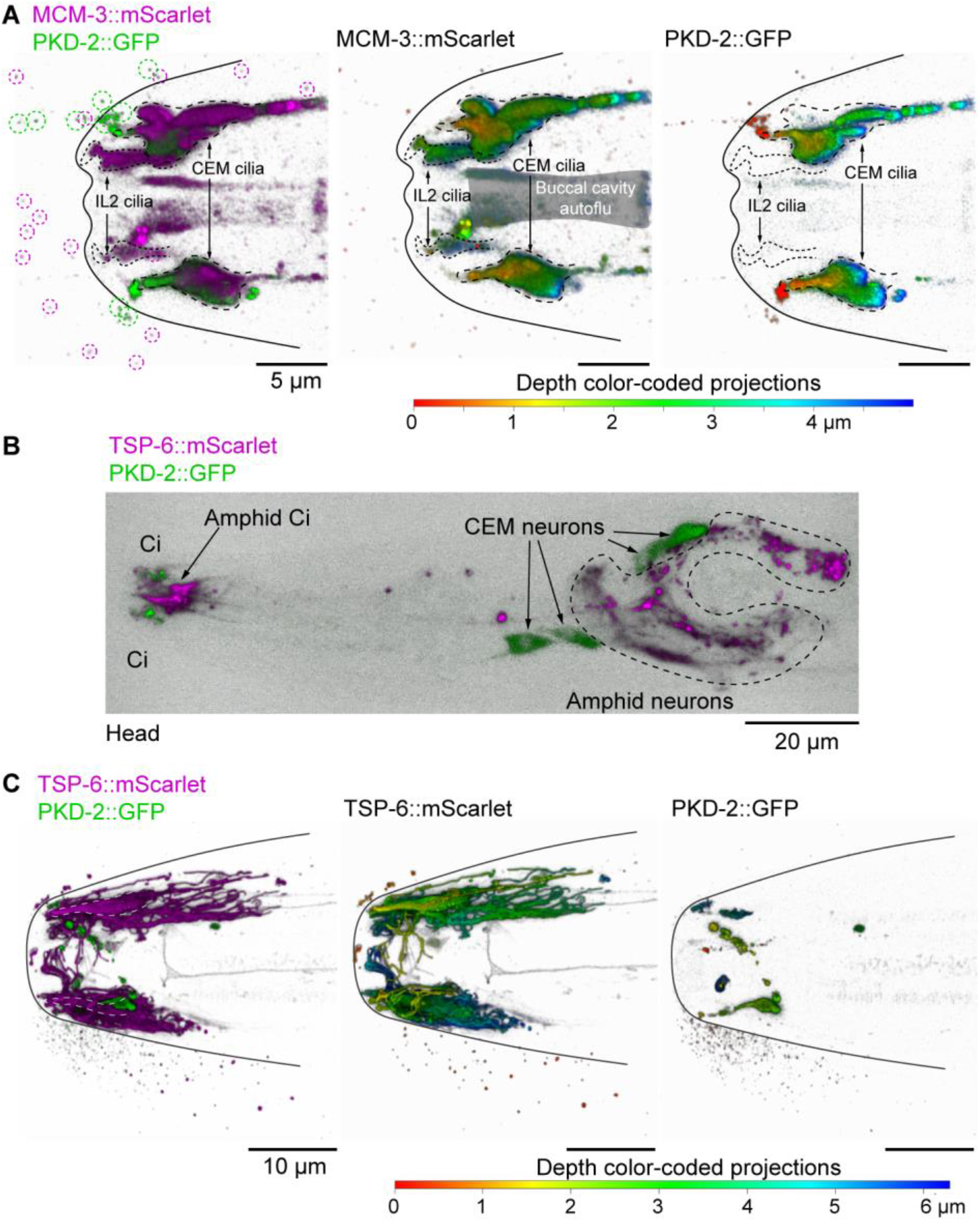
Different neuronal types carry distinct EV cargo; a single neuronal cilium may shed multiple cargoes. (**A**) MCM-3::mScarlet is present in cilia of PKD-2-expressing and IL2 neurons. Circles indicate MCM-3::Scarlet (magenta) and PKD-2::GFP EVs (**B-C**) TSP-6::mScarlet is present in amphid channel and AWA neurons of the head (C) and is released in ciliary EVs into the environment (D). Amphid channel is outline with white dashed line. Merged 3D projections are shown; individual channels are presented as depth color-coded projections with minimal labeling.

Functions of MCM proteins outside of replication are known within Metazoa. For example, a paralog of MCM-3, the MCM2 protein, is required for normal cilia function during *Zebrafish* development [48] and is enriched in sensory ciliated hair cells of the inner ear [49]. Mutations in human *MCM2* are associated with autosomal dominant deafness. Our finding that *mcm-3* is expressed in sensory ciliated EVNs of *C. elegans* suggests a conserved role for MCM components in the physiology of ciliated cells and ciliary EV signaling.

Finally, we also explored EV cargo candidates enriched in the ciliated neurons but specifically excluded from IL2 neurons (Category III, Figure 2F). For that, we examined the expression pattern of TSP-6, an ortholog of human tetraspanin CD9, a marker for human extracellular vesicle isolation [16] (Figure 5B-C). The TSP-6::mScarlet was present in filament-like branches of the chemosensory AWA cilium, rod-like amphid channel cilia of the head, as well as the phasmid cilia of the tail (Figure S5D-E) but was not visibly expressed in EVNs. We observed the TSP-6-carrying EVs being released from a channel of the amphid sensillum (Figure 5C). At the time of this writing TSP-6 was independently discovered as an EV cargo that can also be taken by surrounding amphid glial cell [50], suggesting the role of EVs in the neuron-glia crosstalk. Collectively, these findings open a new field of studying ciliary EVs in diverse ciliary contexts in *C. elegans* and suggests that all cilia may have the potential to shed EVs.

### Males transfer seminal EVs to the hermaphrodite uterus during mating

We also observed that ENPP-1::mScarlet localized to the male reproductive tract. The ENPP-1::mScarlet localization pattern in the secretory cuboidal cells of the vas deferens resembled giant multivesicular bodies (MVB) forming intrusions reminiscent of an ongoing exosome-formation process (Figure 6A, S4D). ENPP-1 localization to the male reproductive tract is consistent with reports of ENPP3 being abundant in the EVs of the epididymal fluid of mammals [51,52]. Thus, we tested whether the ENPP-1 EVs were transferred from males to hermaphrodite partners during mating. We paired an *enpp-1::mScarlet*-expressing male to a non-fluorescent hermaphrodite for copulation (Figure 6B) and observed ENPP-1::mScarlet EV release from the male tail as he backed along the hermaphrodite’s body scanning for the vulva (Movie S2). Upon inserting his copulatory spicules through the vulva, the male transferred seminal fluid containing ENPP-1::mScarlet EVs into the uterus (Figure 6C, Movie S2). We hypothesize that the giant MVB-like structures of the *C. elegans* reproductive tract are an equivalent of the apocrine secretions of the epididymal epithelium of mammals that carry loads of smaller EVs into the seminal fluid [2], ENPP-1 EVs maybe similar to mammalian epididymosomes that are of densities similar to PKD-2::GFP EVs [53]. Overall, these observations indicate that the same EV cargo can function in multiple tissues with different modes of EV biogenesis, i.e. ectocytosed in the form of ciliary EVs or released from MVBs as exosomes.

**Figure 6.**
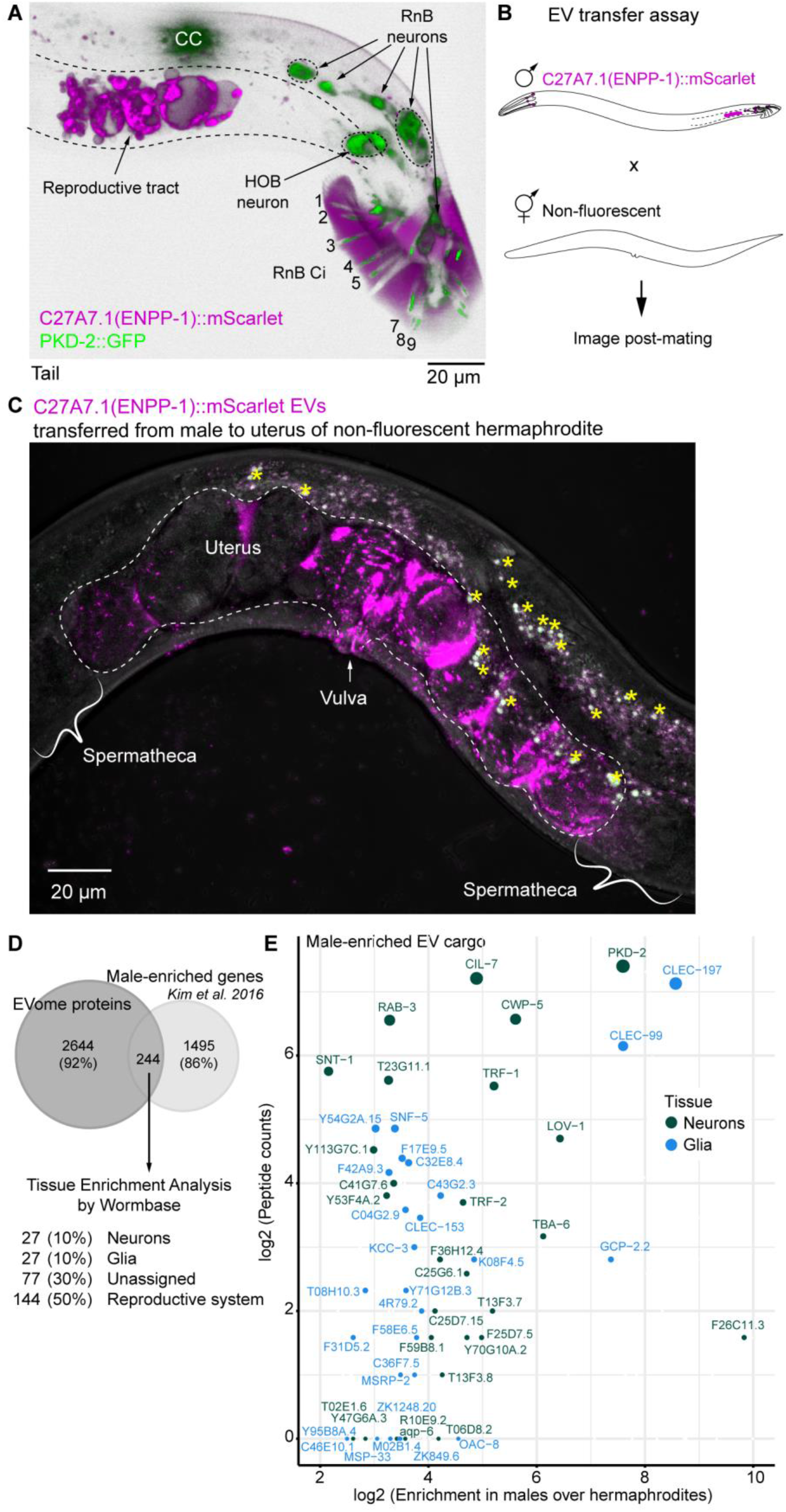
Males transfer seminal EVs to the hermaphrodite uterus during mating. (**A**) ENPP-1::mScarlet in the male reproductive tract. PKD-2::GFP reporter marks male-specific ciliated neurons (RnB and HOB), Ci – cilia, CC-coelomocytes. Merged 3D projection is shown; for individual channels see Figure S4. (**B**) EV transfer experiment included mating ENPP-1::mScarlet males to non-fluorescent hermaphrodites. (**C**) Imaging of the non-fluorescent hermaphrodite after copulation with ENPP-1::mScarlet male revealed ENPP-1 EV presence in the uterus. Yellow asterisks indicate intestinal auto fluorescent droplets. See Movie S2 for real-time imaging of male transferring ENPP-1::mScarlet EVs into the hermaphrodite uterus. (**D**) Venn diagram showing overlap of the identified EV proteome with products of the male-enriched transcripts [54]. (**E**) Scatter plot showing EV cargo predicted to be of either neuronal or glial origin.

Since our dataset contained seminal fluid EVs, we proceeded to compare our dataset with the list of gene products overrepresented in males as identified by bulk transcriptomics [54] (Figure 5D, Table S3). We found 244 male-enriched or male-specific gene products, among which more than 50% were predicted to originate from the male reproductive tract, whereas others were predicted to arise from male-specific neurons or glia (Figure 5E). Thus, our study represents the first report of a male-specific EV proteome that largely expands the list of conserved and species-specific animal EV cargo.

### Data mining tool to identify cell- and tissue- specific EVomes

Our dataset can be mined to predict tissue-specific EV cargo shed into the environment using the same scoring approach we used to identify new ciliary EV cargo. For example, among hypodermis-specific peptides (Figure S6A), we found proteins involved in maintenance of cuticle integrity. Those include CPI-1 inhibitor of cysteine proteinases, which are often used by plants as nematocidal agents [55], and NIT-1 nitrilase, a hydrolase of deaminated glutathione [56], that is a likely byproduct of glutathione-mediated cuticular shedding during molting [57]. We also found GRD-3, a hedgehog-related protein specific to seam cells. Seam cells are known to release exosomes to build alae [58], cuticular ridges running along the *C. elegans* body. Other hypodermis-enriched EV cargo candidates included proteins involved in cholesterol and one-carbon metabolism (SAMS-1, SAMT-1, PMT-1, PMT-2) [59,60], and melanin/tyrosine metabolism (PAH-1, HPD-1, TYR-4) [61–64]. Cuticle composition is regulated in response to different developmental and environmental cues [65–67]. We hypothesize that one way to achieve this would be to deliver matrix-modifying enzymes in the form of EVs to the cuticular extracellular space.

Another example of an EV source tissue that was highly represented in our EVome was the germline (Figure S6B). The germline-specific EV cargo candidates included proteins implicated in formation of extracellular matrix of the eggshell (EGG-1, EGG-2, EGG-6, PERM-4, CBD-1, CPG-1, CPG-2, GLD-1) [65] and RNA metabolism associated with germ granules: Y-box binding proteins, orthologs of YBX1 implicated in sorting RNAs into EVs (CEY-1, CEY-2, CEY-3, CEY-4) [68,69], Argonautes (CSR-1, PPW-1, PPW-2, WAGO-1, WAGO-4) [70], P-granule assembly proteins (CGH-1, PGL-1, PGL-3, DEPS-1, EGO-1, PID-2, PRG-1, ELLI-1, GLH-1, OMA-1, PUF-5) [71], and many other nucleic acid binding proteins, such as RUVB-1, M28.5, all MCM family members (for the full list refer to Table S2). Overall, overrepresentation of nucleic acid binding proteins (Figure 2B, Table S2) in our dataset suggests that environmental EVs could serve as vectors for horizontal transfer of genetic information and regulation of gene expression in trans.

## Discussion

Our work establishes *C. elegans* as a discovery platform for animal EV biology. We obtained a much greater depth of *C. elegans* EV protein identification than in previous studies [72,73] via use of a mixed-stage *C. elegans* population with males and hermaphrodites, enrichment via buoyant density centrifugations, and direct visualization of GFP-labeled EV fractions of interest. The proteogenomic mining approach that we applied to EV cargo candidates lead to the discovery of four novel ciliary EV cargoes. Based on *in vivo* fluorescent protein-tagging and imaging, none of the newly discovered cargoes co-localized to PKD-2 on ciliary EVs, indicating great complexity and heterogeneity of the EV mixture that co-purifies with PKD-2 EVs. Our results show that EVs shed by different neuronal types carry distinct EV cargo and corroborate our previous report that a single neuronal cilium may shed multiple cargoes [13].

Enrichment of our dataset in nucleic acid binding proteins is in agreement with findings of abundant small RNAs carried by EVs of parasitic nematodes [14]. Our work demonstrates that ciliary EVs carry RNA- and DNA- binding proteins, suggesting that cilia may play a role in horizontal transfer of genetic information. Many EV cargo candidates were predicted to originate from the male reproductive tract and thus might play roles in regulation of gene expression [74], fitness of male spermatozoa, male fertility, and sexual interactions [75].

*C. elegans* offers many advantages as a discovery system for identifying EV cargo, mechanisms of EV biogenesis, and EV-based communication pathways, including genetic tractability, *in vivo* imaging and bioassays, and scalability necessary for identification of rare and/or signaling-induced EV cargoes. Our proteogenomic mining strategy can be applied to any cell- or tissue-of-interest using the MyEVome Web App. We envision that our dataset can be exploited and used as an experimental springboard in three different ways: (i) identification of conserved and species-specific drivers of EV-based communication in general, and ciliary EV biogenesis in particular, (ii) discovery of novel bioactivities that have not been associated with extracellular vesicles, (iii) in combination with other identified EV proteomes, understanding of evolutional specialization of EV-based communication within Nematoda clades, including parasites that affect up to a quarter of the human population [76].

## Acknowledgments

We are grateful to Haiyan Zheng for assistance with the mass spectrometry analysis, Gloria Androwski and Helen Ushakov for outstanding technical assistance, Piali Sengupta, Rachel Kaletsky, Coleen Murphy, Julie Claycomb, and Tetsuya Nakamura for discussions, and members of the Boston and UCSF cilia supergroups and Rutgers *C. elegans* community for thought-provoking questions. We also thank WormBase (release WS281), WormAtlas, and WormBook that were used daily during this project.

## Author contributions

Conceptualization: IAN, JW, MMB

Methodology: IAN, JW, JDW, ALC, PT, PS

Investigation: IAN, JW, KMP, JSA, ALC

Visualization: IAN, JW, KMP, JSA, PT

Funding acquisition: MMB, JDW, PS

Project administration: IAN, JW, MMB, PS

Supervision: MMB, PS

Writing – original draft: IAN

Writing – review & editing: IAN, JW, ALC, KMP, JDW, JSA, ARK, PS, MMB

## Data and materials availability

Raw proteomics data generated in this study are available at https://massive.ucsd.edu/ (Accession number – MSV000087943). Generated plasmids and animal strains are available upon request.

## Supplementary Materials

### Materials and Methods

#### Strains

| Name | Genotype | Purpose |
| --- | --- | --- |
| PT3112 | *pha-1(e2123ts) III; him-5(e1490) myls4[pkd-2p::pkd-2::gfp + unc-122p::GFP] V; myEx888 [Pcil-7::cil-7::tagrfp + pBX1]* | EV isolation [13] |
| PT2102 | *pha-1(e2123) III; him-5(e1490) V; myEx686 [klp-6p::gfp::klp-6::3'UTR + pBX1]* | EV cargo validation [77] |
| PT3545 | *pha-1 (e2123ts) III; myls1 [pkd-2p::pkd-2::gfp + unc-122p::GFP] IV; him-5 (e1490) V; myEx944[tsp-6p::tsp-6::mScarlet 20 ng/ul + pBX1 100 ng/ul]* | EV cargo validation |
| PT3562 | *sid-2 (my95[sid-2::mScarlet]) III; myls4[pkd-2p::pkd-2::gfp + unc-122p::GFP] him-5 (e1490) V.* | EV cargo validation |
| PT3617 | *pha-1 (e2123ts) III; myls1 [pkd-2p::pkd-2::gfp + unc-122p::GFP] IV; him-5 (e1490) V; myEx951[mcm-3p::mcm-3::mScarlet 20 ng/ul + pBX1 100 ng/ul]* | EV cargo validation |
| PT3618 | *pha-1 (e2123ts) III; myls1 [pkd-2p::pkd-2::gfp + unc-122p::GFP] IV; him-5 (e1490) V; myEx948[c27a7.1p::c27a7.1::mScarlet 20 ng/ul + pBX1 100 ng/ul]* | EV cargo validation |

#### General maintenance of *C. elegans*

Animals were routinely maintained on 6-cm plates with normal growth medium (NGM) (3 g of NaCl, 2.5 g of peptone, and 20 g of agar per 1 L) seeded with 100 μL *E. coli* strain OP50 [78]. Wild-type strains were propagated at 20°C, whereas temperature sensitive strains carrying *pha-1 (e2123 ts)* alleles were grown at 15°C up until their transgenesis with a cocktail containing a functional *pha-1* allele. After that, the transgenic animals were transferred to a restrictive temperature of 22°C to select for animals carrying the transgene-of-interest [79].

#### Propagation of *C. elegans* for EV enrichment

To initiate the culture, 180 hermaphrodites were transferred to 30 6-cm NGM plates (6 animals per plate) and grown for 2 generations. At day 6, each plate was chunked into quarters and the chunks were transferred into 15-cm plates with high growth medium (HGM) (3 g of NaCl, 2.5 g of peptone, and 20 g of agar per 1 L supplemented with 4 mL of cholesterol stock (5 mg/mL of ethanol), 1 mL 1M CaCl_2_, 1 mL 1M MgSO_4_, and 25 mL 1M potassium phosphate buffer pH 6.0) seeded with a full lawn of *E. coli* OP50 (1.8 mL/plate). Animals were allowed to propagate for 2 more generations until almost all bacteria were consumed, yet animals were not starved.

#### *C. elegans* harvest and EV enrichment procedure

Animals were collected into a shallow dish with the M9 buffer (3 g KH_2_PO_4_, 6 g Na_2_HPO_4_, 5 g NaCl, 1 ml 1 M MgSO_4_ per 1 L of ultrapure water) by inverting the 15-cm plates and their gentle agitation against the surface of the buffer. A total of 200 mL of M9 buffer was used to release worms from 30 HGM plates. The collected worm suspension was centrifuged at 3,000 g for 15 minutes at 15°C in 15 mL conical tubes to pellet the worms and bacteria. The supernatant was transferred to fresh tubes and centrifuged at 10,000 g for 30 minutes 3 times at 4°C (in SS34 fixed angle rotor) to get rid of residual bacteria. Cleared supernatants were then transferred to fresh 12 mL Beckman Coulter round bottom tubes and layered with a 2 mL 36% iodixanol cushion. The cushion was prepared by mixing 3 parts of 60% iodixanol (OptiPrep, Sigma #D1556) and 2 parts of 8% sucrose to achieve final density of 1.2 g/L. EVs were collected onto the cushion via centrifugation in the SW41 Ti swinging bucket rotor at 30,000 rpm (110,000 g average) for 70 minutes at 4°C. EV pellets were collected with a thin glass Pasteur pipette, and further diluted either with M9 buffer to a final density of less than 1.07 g/mL for top loading or with 60% iodixanol to final density of 1.18 g/mL for bottom loading. The gradients were prepared ahead of time by a freeze-thaw method. Specifically, 12%, 16%, 20%, 24%, 28%, and 32% iodixanol solutions were made by mixing solution A (8% sucrose) with solution B (60% iodixanol) so that final osmolarities ranged from 270 to 320 mOsm and densities ranged from 1.08 to 1.20 g/mL. The solutions were then layered in a stepwise manner starting from the heaviest layer and freezing the content at −80°C after each added layer. The gradients were thawed at room temperature 1-2 hours prior to use and remained at 4°C up until loading. Loading was performed using a glass Pasteur pipette. Isopycnic centrifugation was carried out for 16 hours at 30,000 rpm using the SW41 Ti Beckman Coulter rotor at 4°C. Fractions were collected using the BioComp’s Piston Gradient Fractionator. Aliquots were taken from each fraction to measure density, protein concentration, and examine enrichment with PKD-2::GFP-carrying EVs. Density was measured by weighing out 150 μL of the examined solution on an analytical scale to calculate weight per volume ratio for each fraction. Protein concentration was measured using Bicinchoninic Acid (BCA) Protein Assay Kit (Pierce, Thermo Fisher #23225) according to manufacturer’s instructions in the presence of 0.2% SDS. PKD-2::GFP EV enrichment was assessed by scanning through an array of representative droplets from each fraction using a Zeiss LSM 880 confocal microscope and its Airyscan detector. Once the most enriched fractions were identified, total protein was isolated from them via methanol-chloroform extraction. Briefly, four volumes of methanol were added to one volume of the EV-enriched fraction, vortexed vigorously for 10 seconds, followed by addition of one volume of chloroform and repeated intense vortexing, followed by addition of three volumes of ultrapure water and vortexing. Phase separation was completed by centrifugation at 16,000 g in a tabletop centrifuge for 1 minute. The aqueous upper layer was discarded without disturbing the white flaky proteinaceous interphase. Protein was precipitated by adding four volumes of methanol and centrifugation at 16,000 g for 10 minutes. Protein pellets were dried and dissolved in 2x Laemmli buffer (0.125 M Tris HCl pH6.8, 4% SDS, 20% glycerol, 10% 2-mercaptoethanol, 0.004% bromophenol blue) at 90°C for 5 minutes. Resulting proteins were analyzed by denaturing electrophoresis in polyacrylamide gel (SDS-PAGE) or sent for protein identification by mass spectrometry.

#### Transmission electron microscopy

Fractions most enriched with PKD-2::GFP carrying EVs were diluted 10-20 times with M9 buffer and applied to discharged formvar/carbon coated copper grids (200 mesh, Electron Microscopy Sciences # FCF200-Cu). Five microliters of the diluted fraction were dispensed on the grid and allowed on for 30-60 seconds followed by wicking the liquid away from the grid with Whatman filter paper #1. The grid was then transferred through 3 droplets of PBS (5 min in each droplet) to wash away residual sucrose and iodixanol. The excess of PBS was wicked away with the filter paper after each wash. For the unfixed preparation, following PBS washes the grid was exposed to the Nano-W stain (Methylamine Tungstate solution, Nanoprobes #2018) for 1 minute sharp. The stain was removed with the filter paper and the grid was left to dry out completely for 30 minutes at room temperature. Imaging was performed on Philips CM12 electron microscope with AMT-XR11 digital camera. Best results were achieved with imaging the samples on the day of their staining.

#### Liquid chromatography-tandem mass spectrometry (LC-MS-MS)

Samples were allowed to run in an acrylamide gel for a distance of 3-5 cm. Regions-of-interest were excised for further protein processing. Each piece was subjected to reduction with 10 mM dithiothreitol for 30 min at 60°C, alkylation with 20 mM iodoacetamide for 45 min at room temperature in the dark, and digestion with sequencing grade trypsin (Thermo Scientific #90058) overnight at 37°C. Peptides were extracted twice with 5% formic acid, 60% acetonitrile and dried under vacuum.

Samples Light_1 and Heavy_1 were analyzed using Dionex UltiMate 3000 RLSCnano System, ThermoFisher) interfaced with QExactive HF (ThermoFisher). Samples were loaded onto a fused silica trap column Acclaim PepMap 100, 0.075 mm × 200 mm (ThermoFisher). After washing for 5 min at 5 μl/min with 0.1% trifluoroacetic acid, the trap column was brought in-line with an analytical column (nanoEase M/Z Peptide BEH C18 column, 130 Å, 1.7 μm, 0.075 mm × 250 mm, Waters) for LC-MS-MS. Peptides were fractionated at 300 nL/min using a segmented linear gradient of 4-15% solution “B” in solution “A” for 30 min (where A is a polar solvent 0.2% formic acid, and B is an organic solvent of 0.16% formic acid and 80% acetonitrile), 15-25% for 40 min, 25-50% for 44 min, and 50-90% B for 11 min. The column was re-equilibrated with 4% solution “B” in solution “A” for 5 minutes prior to the next run. Cyclic series of full scan acquired in Orbitrap with resolution of 120,000 were followed by MS-MS (HCD relative collision energy 27%) of the 20 most intense ions at resolution 30,000 and dynamic exclusion duration of 20 sec.

For the samples Mixed, Light_2, Heavy_2, Light_3, Heavy_3, Light_4, Heavy_4, Light_5, Heavy_5, Light_6, Light_7, Heavy_6, LC-MSMS were done using Dionex UltiMate 3000 RLSCnano System interfaced with Eclipse Orbitrap tribrid (ThermoFisher). The LC method is the same as described above. For mass spectrometry, the scan sequences began with MS1 spectrum (Orbitrap analysis, resolution 120,000, scan range from M/Z 375–1500, automatic gain control (AGC) target 1E6, maximum injection time 100 ms). The top S (3 sec) and dynamic exclusion of 60 sec were used for selection of parent ions of charge 2-7 for MS-MS. Parent masses were isolated in the quadrupole with an isolation window of 1.2 m/z, AGC target 1E5, and fragmented with higher-energy collisional dissociation with a normalized collision energy of 30%. The fragments were scanned in Orbitrap with resolution of 15,000. The MS-MS scan range were determined by charge state of the parent ion but lower limit was set at 110 amu.

#### Protein identification

The peak list of the LC-MS-MS were generated by Thermo Proteome Discoverer (v. 2.1) into MASCOT Generic Format (MGF) and searched against databases of *C. elegans* (Ensembl), *E. coli* K12 substrain MG1655 (NCBI), and Contaminant Repository for Affinity Purification Mass Spectrometry Data (CRAP) using in house version of X!Tandem GPM (The Global Proteome Machine) Fury [80]. Search parameters were as follows: fragment mass error 20 ppm, parent mass 5 error +/− 7 ppm, fixed modification – carbamidomethylation on cysteine, variable modifications – oxidation on methionine, protease specificity – trypsin (C-terminal of R/K unless followed by P), with 1 miss-cut at preliminary search and 5 miss-cuts during refinement. To validate the protein and peptides, false positive rate (FPR) [81] value was used to decide the log(e) cutoff. Only spectra with log(e) < −2 were included in the final report.

#### Analysis of differential abundance in light vs. heavy fractions

For the differential abundance analysis, the “Mixed” dataset was omitted as these EVs did not resolve in two populations. For the rest of the samples Normalized Spectral Abundance Factors (NSAFs) were calculated from uniquely mapped spectral counts. Batch corrections were applied using the R package msmsEDA. For differential expression analysis, we considered only proteins that were identified in more than one replicate for both heavy and light subpopulations. For these proteins, missing values were imputed using the QRILIC method as implemented in the R package DEP [82]. Differential expression analysis was performed using the R package limma [83], using the trend and robust hyperparameter estimation settings. P-values were adjusted using the Benjamini-Hochberg procedure. Proteins were considered differentially expressed between the heavy and light EVs if they had an absolute log2 fold changes greater or equal to 1.5 and an adjusted p-value < 0.05.

#### Gene ontology and InterPro enrichment

KEGG enrichment was performed using clusterProfiler R package. KEGG enrichment in the heavy EVs included proteins that were upregulated relative to the light EVs and those proteins only detected in the heavy EVs. A similar procedure was performed for light EVs. The background for KEGG enrichments of differentially abundant proteins included only proteins detected via LC-MS-MS. InterPro domain enrichment was performed using a hypergeometric test in a custom R script, with p-values corrected using the Benjamini-Hochberg procedure.

#### Coupling the mass spectrometry data with single cell transcriptomics of *C. elegans*

EV cargo were assigned scores derived from single-cell transcriptomics datasets [37]. Enrichment in ciliated neurons was calculated as the ratio of transcript abundance (transcript per million, TPM) in ciliated neurons over mean transcript abundance across all the tissues. Enrichment in IL2 neurons was calculated as the ratio of transcript abundance in IL2 neurons over its mean abundance in all neurons. Identity of the IL2 neurons was established as Cholinergic clusters 15 from [37] based on similarities of their transcriptional profiles to the list of transcripts enriched in the EV-releasing neurons identified in [10]. The similarity was determined by hierarchical clustering analysis of the fold change enrichment values of EVN-specific transcripts from [10] to the fold change enrichment of respective cholinergic neurons transcripts over all identified cholinergic neuronal clusters of *C. elegans*. The values that were used for generation of the dendrogram on Figure 2E are indicated in Supplemental table S3.

#### Transgenesis for EV cargo validation

Pieces of genomic DNA were cloned into the vector backbone encoding the C-terminally linked flexible linker (amino acid sequence is GGGSGGGSGGGSGGG) fused to the mScarlet coding sequence and *unc-54* 3’UTR using the following primers (annealing regions only are indicated to orient the reader on which genomic piece was cloned):

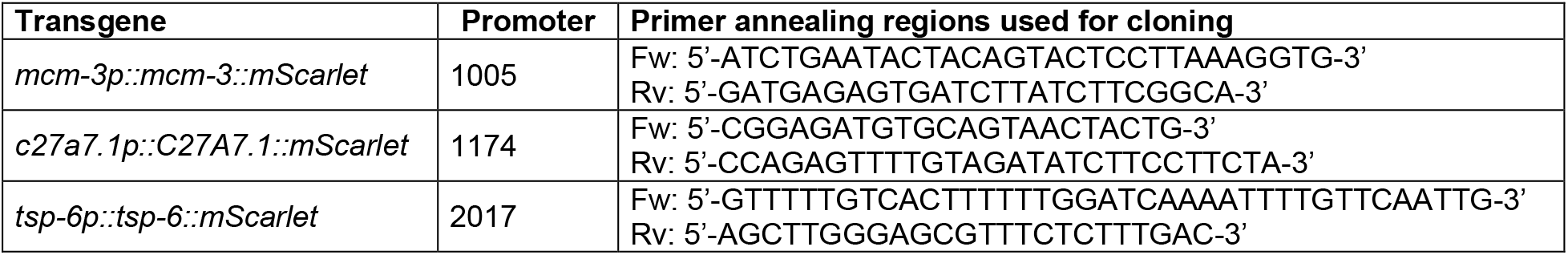

All the transgenes were fully sequenced prior to transgenesis. Respective plasmids were injected with a co-injection selective marker *pha-1(+)* into *pha-1 (e2123 temperature sensitive) III; him-5 V* strain. Further maintenance of the transgenic worms at 22°C ensured selection for and retention of the extrachromosomal transgenic arrays.

The *sid-2::mScarlet* reporter was generated using CRISPR/Cas9-mediatead editing as described in [84] using a guide RNA (5’-CCTTCGCTACATTGGAAAGC-3’) and a double stranded DNA donor template containing a sequence encoding the flexible linker of 15 amino acids (GGGGS)_3_ fused to the mScarlet coding sequence (Figure S3A). The mScarlet coding sequence was intron-less to not interfere with native splicing regulation within the *dyf-2* gene.

#### Mounting animals for super resolution live animal imaging

Animals were anesthetized with 10 mM levamisole solution in the M9 buffer. To completely immobilize the worm, we used thin agarose pads, prepared from 10% melted agarose (Sigma #A9539) and ultrapure water. Melting of the 10% agarose was achieved by pulse heating in a microwave in 3 second intervals for a total of 1 minute or until dry agarose pieces were no longer visible. Once melted, the 10% agarose was incubated at 95°C for about 10 minutes to allow air bubbles to escape. The melted agarose was dispensed onto glass slides in 100 μL droplets and the droplets were immediately pressed with a second glass slide to make a thin gel pad. The agarose pads were made in batches and stored in a sealed glass slide container at room temperature for up to a week. Animals were mounted by their placement in 1 μL droplet of 10 mM levamisole dispensed on a 18 × 18 cm coverslip, followed by flipping of the coverslip onto the center of the agarose pad. Animals were imaged within 30 minutes of their mounting.

#### Super-resolution microscopy

Super-resolution imaging was performed on a Zeiss LSM880 confocal system equipped with Airyscan Super-Resolution Detector with Fast Module, 7 Single Photon Lasers (405, 458, 488, 514, 561, 594, 633 nm), Motorized X-Y Stage with Z-Piezo, T-PMT. Planes of captured Z-stacks were distanced 0.185 μm for optimal 3D rendering. Image analysis and processing was performed using Zen Blue 2.0 and Zen Black 2.0.

#### Live imaging of EV release during mating

To visualize EV release during mating three *unc-51* hermaphrodites were placed with ten males of the PT3618 strain in the center of an unseeded NGM plate, some bacteria were transferred on the pick to feed the hermaphrodites. The piece of agar with the placed worms (approximately 24 × 40 mm) was transferred onto a 24 × 60 coverslip with the worm side facing the coverslip. Then the agar chunk was covered with a 28 × 48 mm saran wrap, with a 10 mm- diameter hole in the center, so that the spot with hermaphrodites was exposed to air through the window. This covering method drove the males toward the center with the hermaphrodites. Mating started within 15 minutes. Imaging was performed using 40x objective.

**Figure S1.**
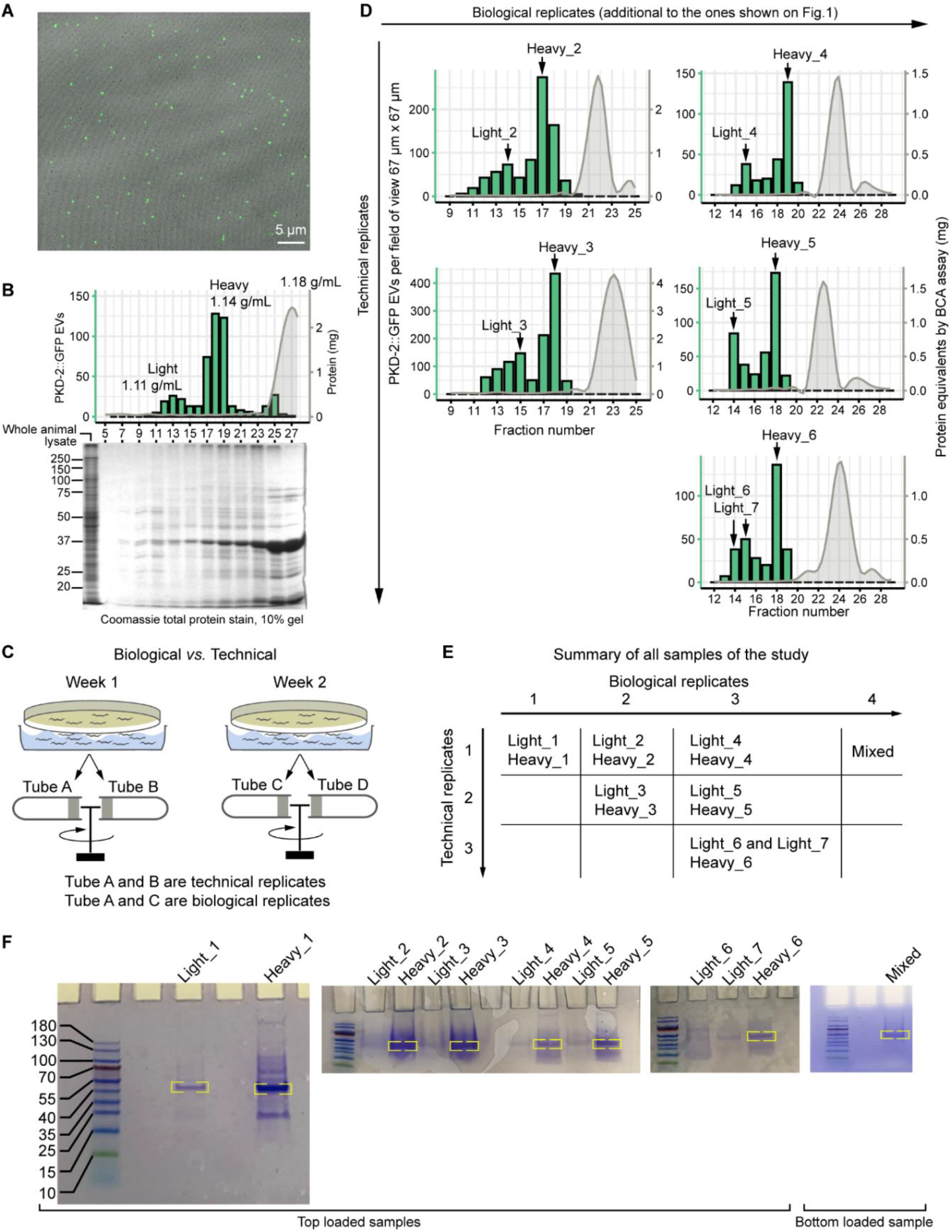
Characterization of the EV preparations prior to the mass spectrometry analysis. (**A**) Representative image of the PKD-2::GFP enriched fraction acquired using Zeiss LSM880 microscope and the Airyscan detection system. (**B**) Analysis of protein content across fractions collected from the isosmotic gradients after buoyant density centrifugation of the crude mixture of the *C. elegans* and *E. coli* EVs. Combination graph shows a profile of the analyzed EV preparation. Image of the Coomassie stained gel shows more complex protein composition of the PKD-2::GFP enriched fractions as compared to the fraction of 1.18 g/mL density enriched in most protein. Approximately 40 μg of protein was loaded into each well. (**C**) Biological replicates are represented by EVs harvested on different weeks, technical replicates are represented by EVs that were collected in one prep but equilibrated for their density in different gradients and fractionated into different tubes. (**D**) Fraction profiles of EV preparations for 2 biological replicates (additional to the ones depicted of Figure 1C), each with 2 and 3 technical replicates, respectively. Labeling indicates names of PKD-2 EV enriched fractions that were further analyzed by mass spectrometry. (**E**) Summary of technical and biological replicates of the study. (**F**) Images of stained gels prior to mass spectrometry analysis. Yellow brackets indicate bands that likely represent bacterial outer membrane proteins, and thus were excised from the gel and were not included into protein identification analysis. This approach enriched the remaining proteins further for the EV cargo of *C. elegans*.

**Figure S2.**
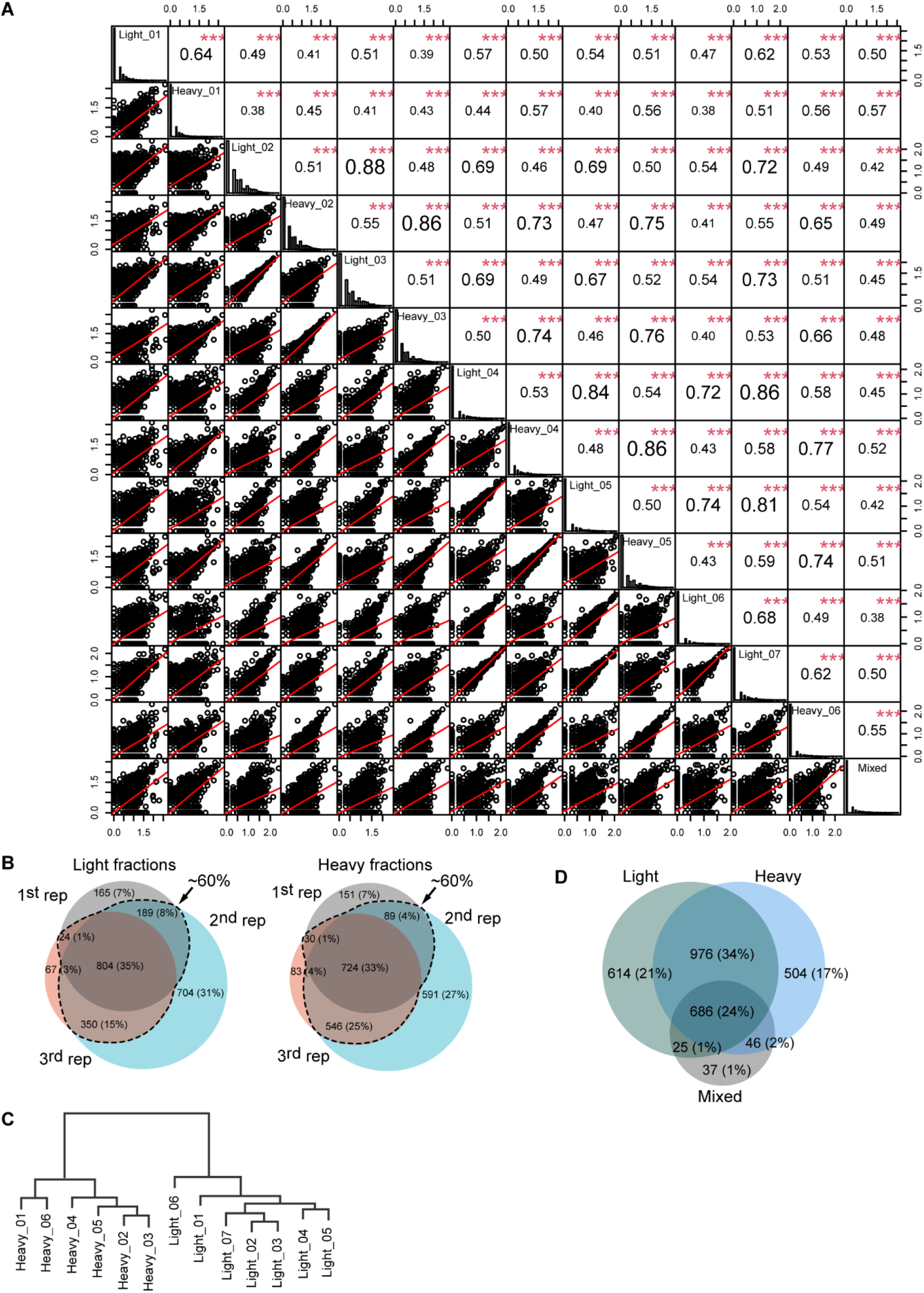
Comparison of the identified protein datasets across technical and biological replicates showed high reproducibility of the EV preparation. (**A**) Sample pairwise comparisons were made using Spearman correlation analysis. Scatter plots at the bottom left corner compare log_10_(Peptide count+1) values in pairwise manner for all samples of the study; Spearman correlation coefficients for each comparison are at the upper right corner; histograms on the diagonal line show frequencies of log_10_(Peptide count+1) values for each sample [85]. (**B**) Venn diagrams showing overlap in the identified protein datasets of the light and heavy PKD-2::GFP EV subtypes obtained from 3 different biological replicates. (**C**) The cladogram produced by hierarchical clustering of the batch-corrected normalized spectral abundance factors (NSAFs) segregates all the light EV samples from all the heavy ones. (**D**) Light and heavy fractions shared 58% common proteins.

**Figure S3.**
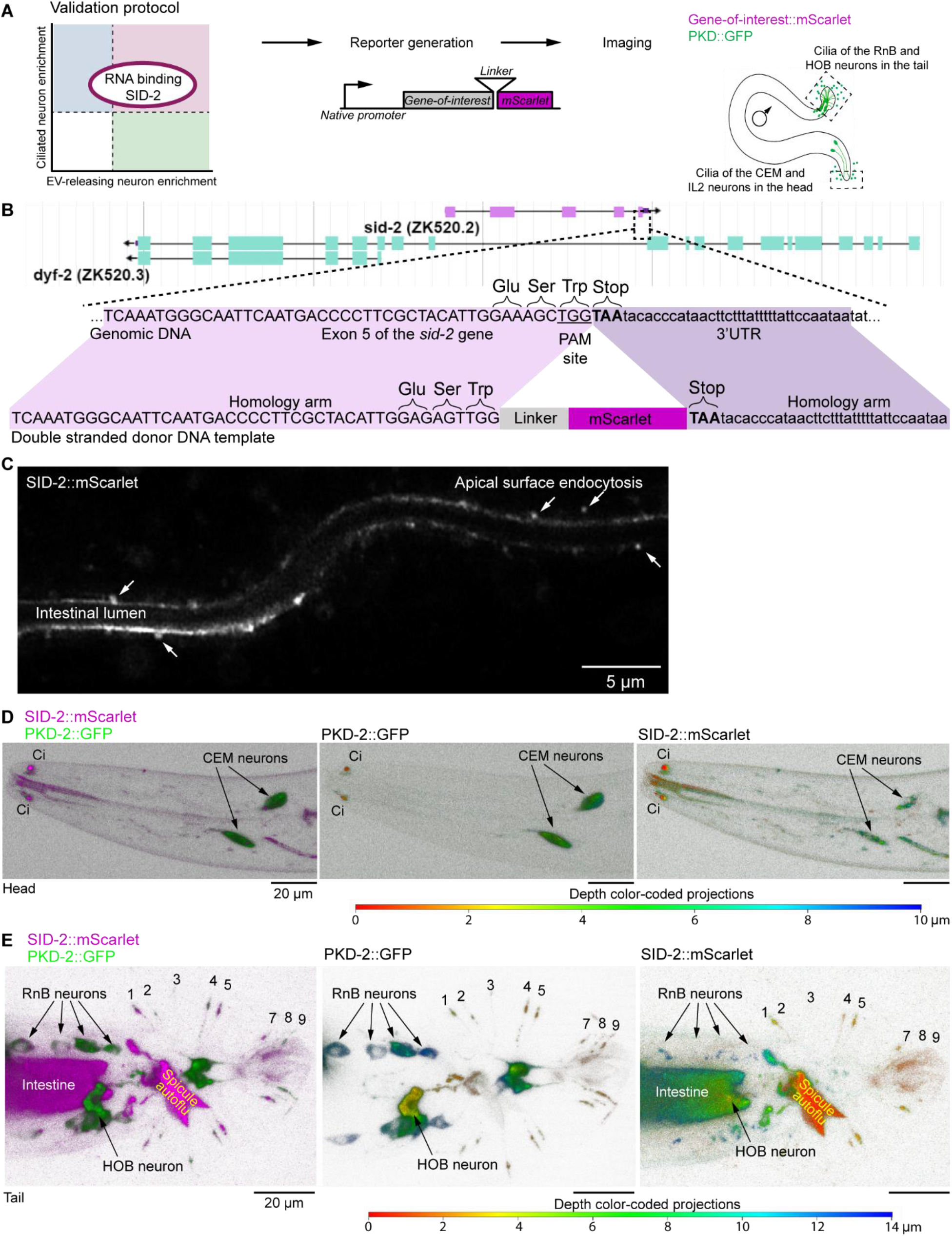
SID-2 nucleic acid binding transmembrane protein is specifically enriched in the male-specific ciliated sensory neurons (CEMs, RnBs, and HOB). (**A**) Validation protocol for putative ciliary EV cargo. (**B**) CRISPR/Cas-9-mediated genome editing of the *sid-2* genomic locus. The *sid-2* gene is situated within a long intron of the *dyf-2* gene encoding an ortholog of human ciliary protein WDR19. Editing was performed to generate the SID-2 gene product fused with mScarlet via a 15 amino acid long flexible linker. Two silent mutations were introduced to the protospacer region encoding -Glu-Ser- residues in order to avoid multiple cuts by the Cas9 protein. PAM stands for “<u>**p**</u>rotospacer <u>**a**</u>djacent <u>**m**</u>otif” and indicates the targeted site for cleavage by Cas9. (**C**) Sagittal image through the intestinal lumen showing enrichment of SID-2::mScarlet on the apical surface of the intestinal cells. Ongoing endocytosis of SID-2 is labeled with white arrows. (**D-E**) 3D projections showing presence of the SID-2::mScarlet reporter in the cilia and cell bodies of the male-specific sensory neurons in the head (D), and in the tail (E). Merged images are shown in 2 colors; individual channels are presented as depth color-coded projections where colocalized presence is indicated by the same color (such as SID-2 and PKD-2 presence within the CEM neurons in (C) is indicated with shades of green and blue which corresponds to ~6 μm distance deep into the acquired Z-stack). Male tail rays housing PKD-2-positive RnB neurons are labeled with numbers. In all male-specific neurons (CEMs, RnBs, and HOB), the SID-2::mScarlet signal was observed in neuronal cell bodies as a few puncta and was mostly accumulated in ciliary regions. SID-2::mScarlet was also observed in two unidentified cells whose processes converge near the hook structure of the male tail.

**Figure S4.**
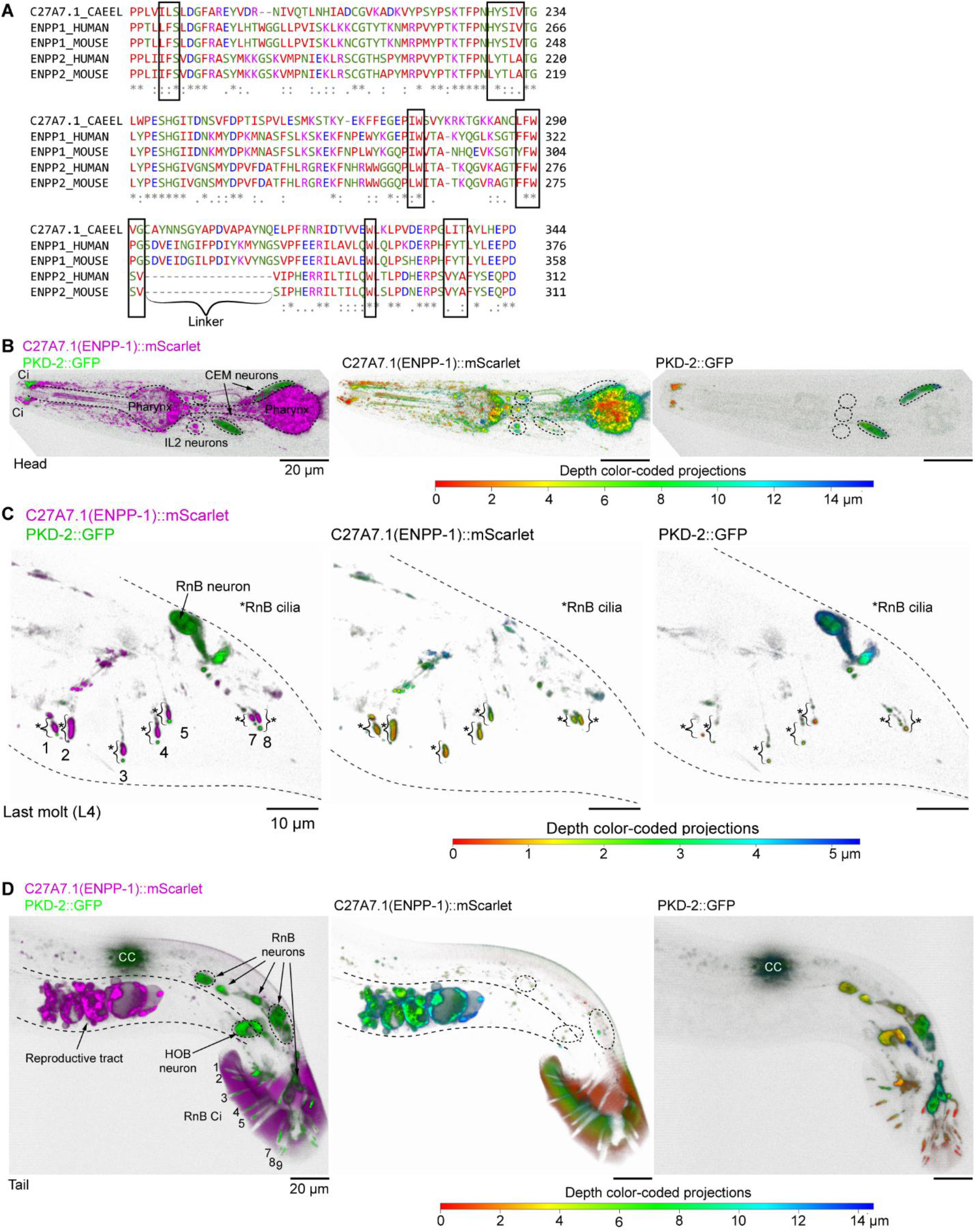
Ortholog of human ENPP1 (C27A7.1) is specifically expressed in the EV-releasing ciliated sensory neurons (IL2s, CEMs, RnBs, and HOB) and male reproductive tract of *C. elegans*. (**A**) Alignment of the phosphodiesterase domain of C27A7.1 to ENPP1 (nucleotide-specific) and ENPP2 (lysophospholipid-specific hydrolase) of human and mouse suggests that C27A7.1 is a nucleotide-specific hydrolase. Black boxes indicate conserved regions forming substrate-binding pockets. Most similarity is observed between C27A7.1 and ENPP1. Figure bracket indicates a linker region of ENPP1 critical for its nucleotide substrate recognition; mutagenic studies show that loss of this linker grants ENPP1 the ability to hydrolase lysophospholipids, yet at much lower efficiency than ENPP2. (**B-D**) 3D projections showing expression pattern of C27A7.1::mScarlet. Merged images are shown in 2 colors; individual channels are presented as depth color-coded projections. C27A7.1 is expressed in pharynx and EV-releasing neurons (EVNs) of the head (CEMs and IL2) (B), and tail (C) as well as in the EVNs of the tail, and the male reproductive tract (D). CC indicates a pair of coelomocytes labeled with GFP as a co-transformation marker of the PKD-2::GFP reporter that remains visible in hermaphrodites. C27A7.1::mScarlet signal also covers the male tail fan indicating that C27A7.1-carrying EVs are adherent to the fan cuticle (D).

**Figure S5.**
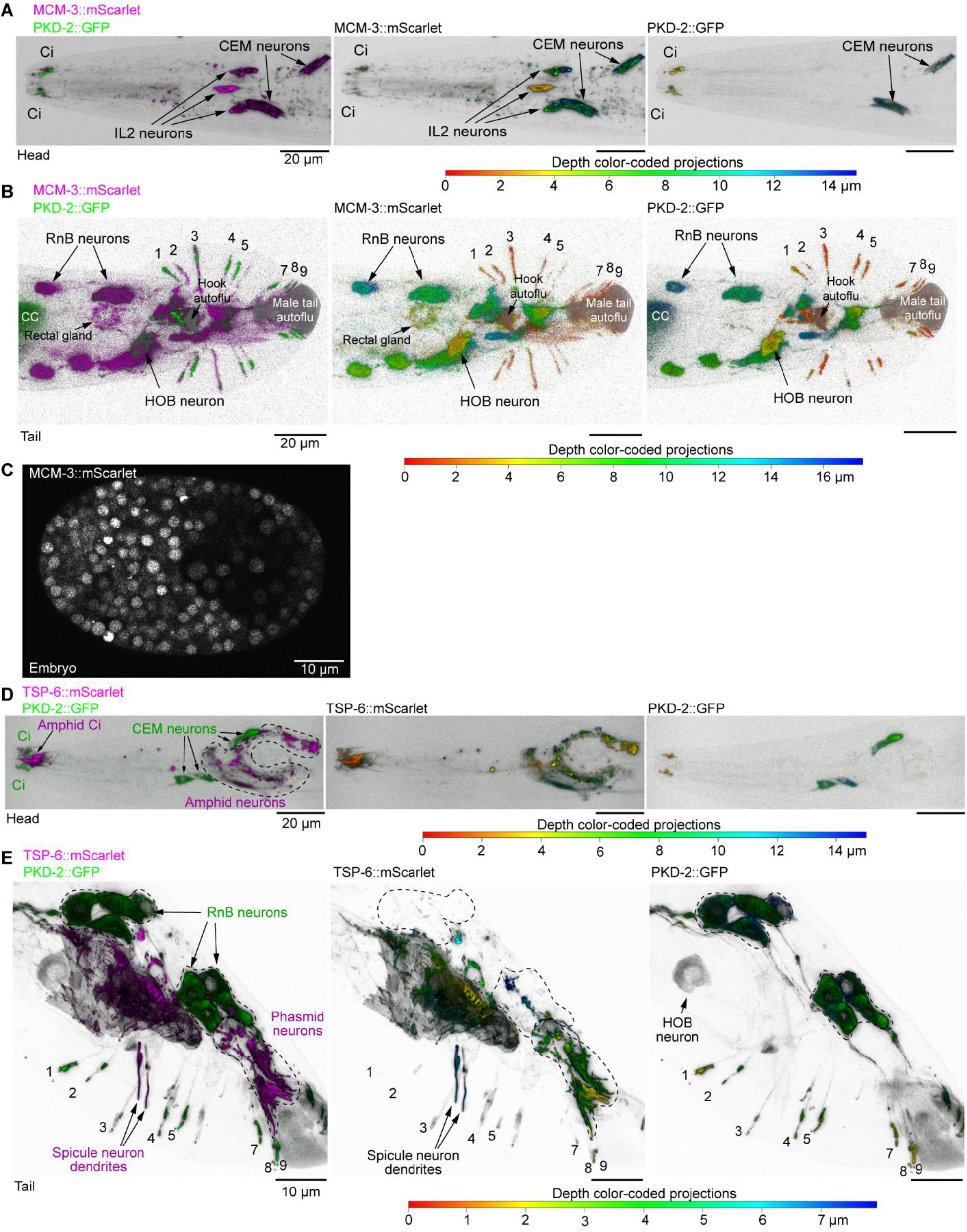
Different neuronal types carry distinct EV cargo; a single neuronal cilium may shed multiple cargoes. (**A-C**) MCM-3 subunit of the replication origin licensing complex is specifically enriched in the EV-releasing ciliated sensory neurons (IL2s, CEMs, RnBs, and HOB). 3D projections showing presence of the MCM-3::mScarlet in the cilia and cell bodies of the EV-releasing sensory neurons in the head – IL2s and CEMs (A), and in the tail – RnBs and HOB (B). Merged images are shown in 2 colors; individual channels are presented as depth color-coded projections. Areas of intense inherent autofluorescence are indicated – the hook structure and the very posterior part of the male tail (B). The MCM-3::mScarlet reporter also shows weak presence in the rectal gland. This identification is based on its similarity to an image in the WormAtlas describing anatomical structures of male proctodeum [86]. (**C**) Sagittal image through the *C. elegans* embryo showed nuclear localization of MCM-3::mScarlet in the actively dividing cells. (**D-E**) Ortholog of human CD-9 (TSP-6) is expressed in the amphid and phasmid ciliated neurons, but not in the PKD-2-expressing EV-releasing neurons. 3D projections showing expression pattern of TSP-6::mScarlet at the fourth larval stage. (**D**) Head region, Ci stands for cilia. (**E**) Tail region. TSP-6::mScarlet is expressed in the spicules and other unidentified structures in the mail tail. Merged images are shown in 2 colors; individual channels are presented as depth color-coded projections. PKD-2::GFP served as a marker of the CEM, RnB, and HOB neurons. RnB neurons are labeled with numbers. CC indicates a pair of coelomocytes labeled with GFP as a co-transformation marker. Cilia of the RnB neurons are indicated with numbers.

**Figure S6.**
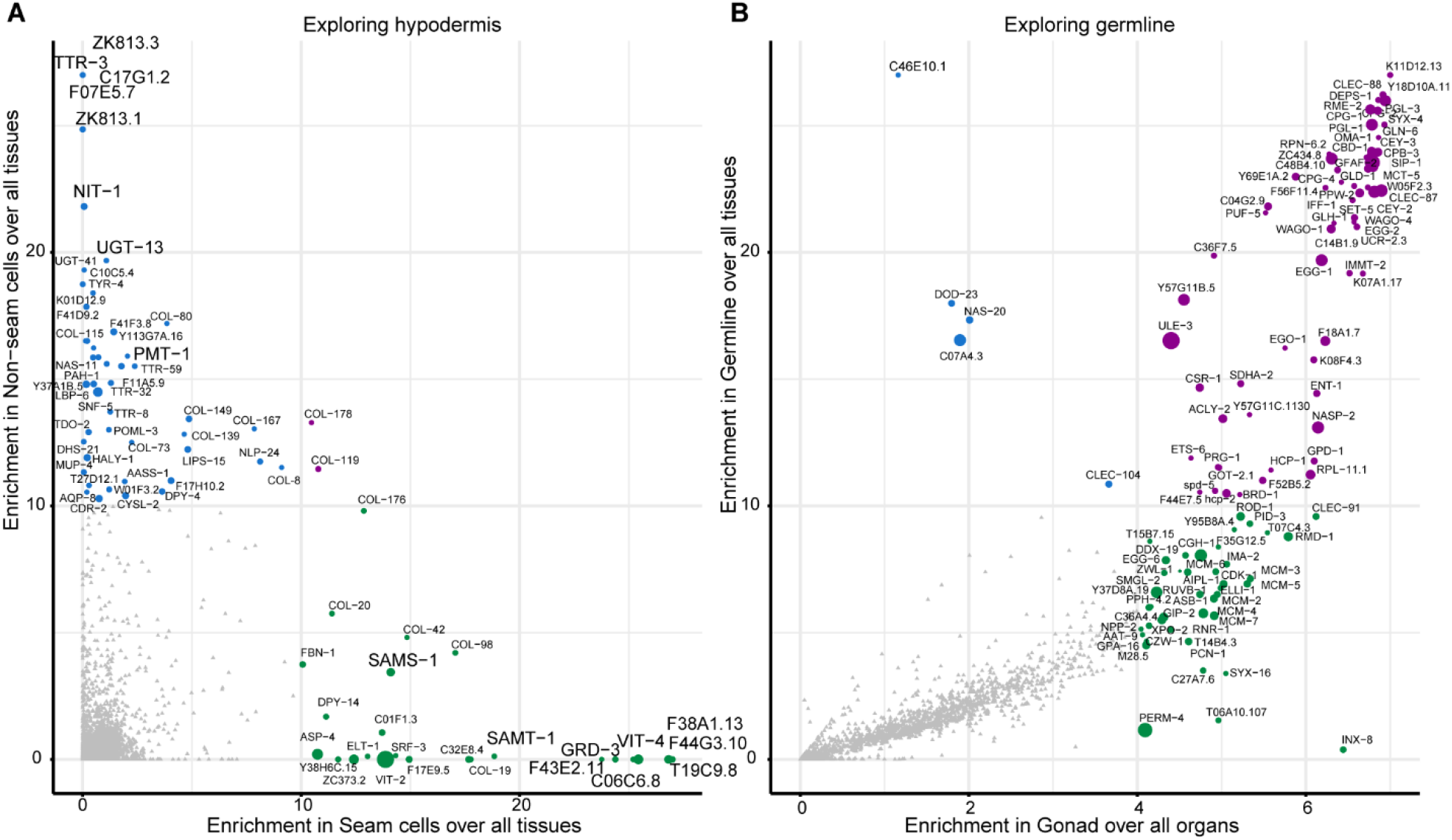
Examples of enrichment plots with tissue-enrichment scores for identified EV cargo candidates. Prediction of EV cargo originating from the hypodermis (**A**) and germline (**B**).

**Movie S1.**

**ENPP-1::mScarlet EV release from the IL2 cilia into the environment.** During the time lapse focus changes from the cilium to EV plane. EVs are connected to each other in a tandem manner and remain attached to the cilium during the course of the video.

**Movie S2**

**ENPP-1::mScarlet EV release from the male tail during mating to a hermaphrodite. The footage shows two males mating with one hermaphrodite.** Male #1 releases ENPP-1::mScarlet EVs while backing along the body of the hermaphrodite scanning for vulva, whereas Male #2 releases ENPP-1::mScarlet EVs during sperm transfer.

**Data S1. (separate .xls file)**

**Lists of identified peptides and proteins**. (**A**) Metadata on the overall numbers of peptides and identified proteins from *C. elegans* and *E. coli*. (**B**) Unique peptides with matched proteins. (**C**) Non-unique peptides matched with corresponding protein sets. (**D**) Summary of spectral counts of unique proteins for each gene product. Functions of the genes and human orthologs are indicated. (**E**) Normalized NSAF values for each identified gene product.

**Data S2. (separate .xls file)**

**Differential abundance and enrichment analyses.** (**A-D**) Differential abundance analyses of the light and heavy EV proteomes together with KEGG pathway and InterPro protein domain enrichment analyses. (**E**) List of ESCRT proteins identified in the EV proteome. (**F**) List of nucleic acid binding proteins identified in the EV proteome.

**Data S3. (separate .xls file)**

**Mining of the EV proteome**. (**A**) Overlap of the EVome with the EVN-specific transcriptomes used to produce Figure 2C. (**B**) IL2 identification table contains relative enrichment values for cholinergic clusters from Cao et al. (2017). (**C**) EV cargo scores based on their absolute enrichment in ciliated sensory neurons and relative enrichment in the IL2 neurons of hermaphroditic L2 populations; these scores were used to generate the plot on Figure 2F. (**D**) Overlap of the identified EV proteome with the male-specific transcriptome from [54].

